## Supplemental Material for "Cryo-EM structure of the mature and infective Mayaro virus at 4.4 Å resolution reveals new features of arthritogenic alphaviruses"

**SUPLEMENTARY MATERIAL**

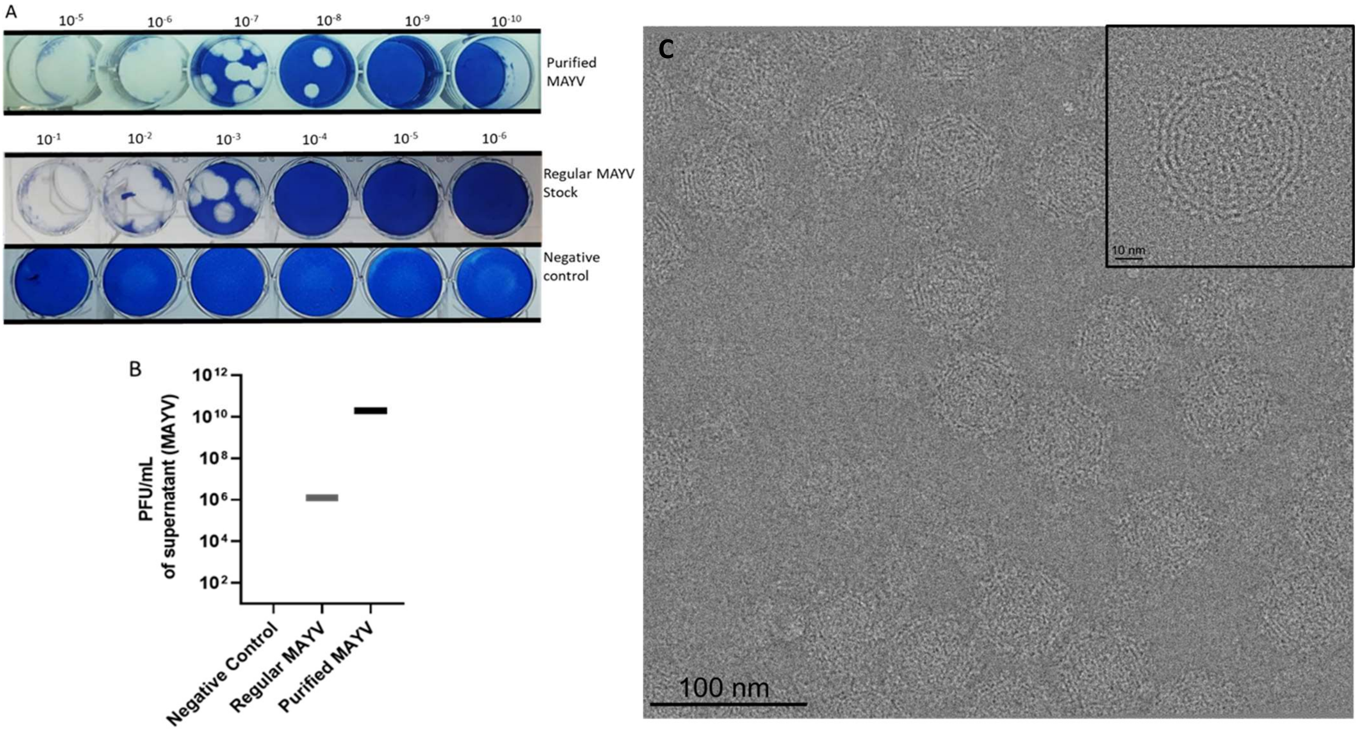

**Figure S1. Purified MAYV stocks used in Cryo-EM experiments are viable and infective. (A)** Representative pictures of plaque assays performed in Vero CCL81 cells to assess infective MAYV load in stock samples. Viable Vero CCL81 cells were stained in Methylene Blue 1% w/v. MAYV replication leads to cell death, creating transparent lysis plaques in the blue Vero CCL81 monolayer. **(B)** Viral load of stocks used in Cryo-EM experiments, before and after the purification process. (C) Representative raw micrograph of MAYV Cryo-EM data collection. Inset presents an individual picked particle used in 3D reconstruction.

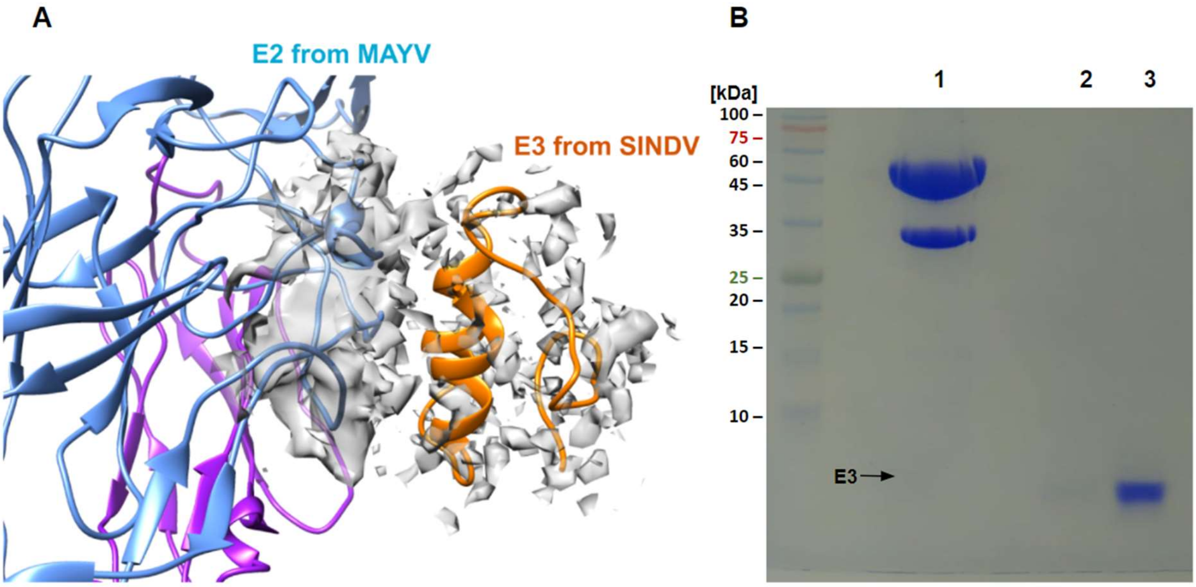

**Figure S2. Absence of the E3 protein indicates that purified MAYV samples are composed of mature virions.** (**A**) Superposition of the MAYV electron density with the SINDV particle cryo-EM structure (entry 6IMM). The SIDNV E3 protein is shown in orange. MAYV electron density is shown in grey contoured at 2.0 sigma-level. (**B**) Analysis of purified MAYV in 15% SDS-PAGE gel, stained with Coomassie brilliant blue. Lane 1 is carried with 30 µg of denatured purified MAYV and shows no visible bands compatible with E3 protein size (7 kDa). Lanes 2 and 3 are carried with 1 µg and 9 µg of a 5 kDa synthetic peptide, used as a positive control.

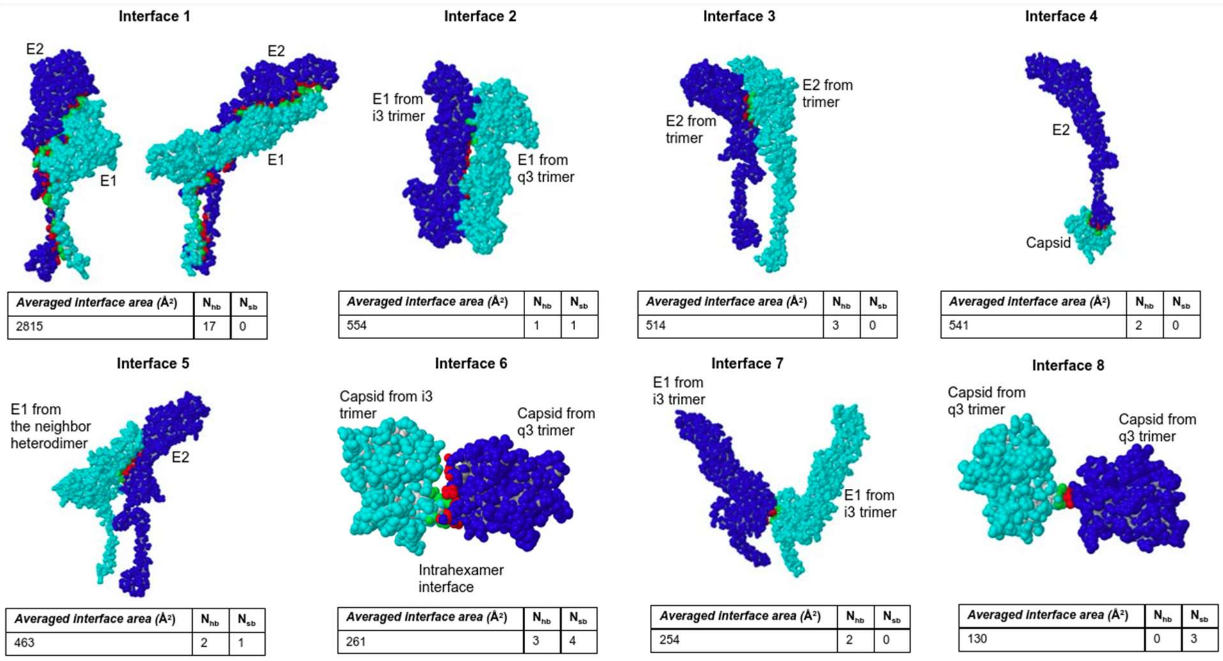

**Figure S3. List of the main interfaces among MAYV structural proteins of the asymmetric unit.** The interfaces were identified using the PDBePISA server (https://www.ebi.ac.uk/pdbe/prot_int/pistart.html). Only interfaces with area larger than 100 Å were considered. The structures are rendered as spacefill and the interaction partners are colored in blue or cyan. Interaction residues are colored in red or green. $N_{hb}$ and $N_{sb}$ represent the average number of hydrogen bonds and salt-bridges in each interface, respectively.

### A - Multiple Sequence Alignment of the Alphaviral Capsid

```
        1        10        20        30        40
MAYV    MDFLPTQVFYGRRWR.PRM.PPR..PW....RPRPPTI....QRPDQQARQMQQLIAAVS
CHIKV   MEFIPTQTFYNRRYQ.PRPWTPR..PTIQVIRPRPR........PQRQAGQLAQLISAVN
SFV     MNYIPTQTFYGRRWR.PRP.AAR..PWPLQATPVAPVV......PDFQAQQMQQLISAVN
RRV     MNYIPTQTFYGRRWR.PRP.AFR..PWQVSMQPTPTMVTPMLQAPDLQAQQMQQLISAVS
SINDV   MN....RGFFNMLGRRPFPAPTA..MW....RPRRRRQAAPMPARNGLASQIQQLTTAVS
VEEV    FPFQPMYPMQPMPYRNPFAAPRR..PWFPRTDP.............FLAMQVQELTRSMA
EEEV    LNYPPMAPINPMAYRDPNP.PRQVAPF....RP.............PLAAQIEDLRRSIA
```

```
        50        60        70        80        90
MAYV    TLALRQ.NAAAPQRGRKKQPRRKKPKPQP......EKPKKQEQKPKQKKTP.........
CHIKV   KLTMRA....VPQQKPRRNRKNKKQKQKQ......QAPQNNTNQKKQPPKKKPAQ.....
SFV     ALTMRQ.NAIAPARPPKPKKKKTTKPKPK......TQPKKINGKTQQQKKKDKQAD....
RRV     ALTTKQ.NVKAP.KGQRQKKQQKPKEKKE......NQKKKPTQKKKQQQKPKPQA.....
SINDV   ALVIGQATRPQPPRPRPPPRQKKQAPKQP......PKPKKPKTQEKKKKQP.........
VEEV    NLTFKQ.RRDAPPEGPSAKKPKKEASQKQKGGGQGKKKKNQGKKKAKTGPPNPKAQNGNK
EEEV    NLTLKQ.RAPNPPAGPPAKRKK.............PAPSLSLETKKKRPPPPAKKQ....
```

```
              100       110       120       130       140
MAYV    ...KKKPGRRERMCMKIEHDCIFEVKHE.GKVTGYACLVGDKVMKPAHVPGVIDNIDLAR
CHIKV   ..KKKKPGRRERMCMKIENDCIFEVKHE.GKVTGYACLVGDKVMKPAHVKGTIDNADLAK
SFV     .KKKKKPGKRERMCMKIENDCIFEVKHE.GKVTGYACLVGDKVMKPAHVKGVIDNADLAK
RRV     ..KKKKPGRRERMCMKIENDCIFEVKLD.GKVTGYACLVGDKVMKPAHVKGTIDNPDLAK
SINDV   ..AKPKPGKRQRMALKLEADRLFDVKNEDGDVIGHALAMEGKVMKPLHVKGTIDHPVLSK
VEEV    KKTNKKPGKRQRMVMKLESDKTFPIMLE.GKINGYACVVGGKLFRPMHVEGKIDNDVLAA
EEEV    .KRKPKPGKRQRMCMKLESDKTFPIMLN.GQVNGYACVVGGRVFKPLHVEGRIDNEQLAA
```

```
      150       160       170       180       190       200
MAYV    LSYKKSSKYDLECAQIPVAMKSDASKYTHEKPEGHYNWHYGAVQYTGGRFTVPTGVGKPG
CHIKV   LAFKRSSKYDLECAQIPVHMKSDASKFTHEKPEGYYNWHHGAVQYSGGRFTIPTGAGKPG
SFV     LAFKKSSKYDLECAQIPVHMRSDASKYTHEKPEGHYNWHHGAVQYSGGRFTIPTGAGKPG
RRV     LTYKKSSKYDLECAQIPVHMKSDASKYTHEKPEGHYNWHHGAVQYSGGRFTIPTGAGKPG
SINDV   LKFTKSSAYDMEFAQLPVNMRSEAFTYTSEHPEGFYNWHHGAVQYSGGRFTIPRGVGGRG
VEEV    LKTKKASKYDLEYADVPQNMRADTFKYTHEKPQGYYSWHHGAVQYENGRFTVPKGVGAKG
EEEV    IKLKKASIYDLEYGDVPQCMKSDTLQYTSDKPPGFYNWHHGAVQYENNRFTVPRGVGGKG
```

```
      210       220       230       240       250
MAYV    DSGRPIFDNKGRVVAIVLGGANEGARTALSVVTWN.KDMVTKITPEGTEEW
CHIKV   DSGRPIFDNKGRVVAIVLGGANEGARTALSVVTWN.KDIVTKITPEGAEEW
SFV     DSGRPIFDNKGRVVAIVLGGANEGSRTALSVVTWN.KDMVTRVTPEGSEEW
RRV     DSGRPIFDNKGRVVAIVLGGANEGARTALSVVTWT.KDMVTRVTPEGTEEW
SINDV   DSGRPIMDNSGRVVAIVLGGADEGTRTALSVVTWNSKGKTIKTTPEGTEEW
VEEV    DSGRPILDNQGRVVAIVLGGVNEGSRTALSVVMWNEKGVTVKYTPENCEQW
EEEV    DSGRPILDNKGRVVAIVQGGVNEGSRTALSVVTWNQKGVTVKDTPEGSEPW
```

### B - Multiple Sequence Alignment of the Alphaviral E1

```
       1        10        20        30        40        50        60        70        80
MAYV   YEHTAVIPNQVGFPYKAHVAREGYSPLTLQMQVVETSLEPTLNLEYITCDYKTKVPSPYVKCCGTAECRTQDKPEYKCAV
CHIKV  YEHVTVIPNTVGVPYKTLVNRPGYSPMVLEMELLSVTLEPTLSLDYITCEYKTVIPSPYVKCCGTAECKDKNLPDYSCKV
SFV    YEHSTVMPNVVGFPYKAHIERPGYSPLTLQMQVVETSLEPTLNLEYITCEYKTVVPSPYVKCCGASECSTKEKPDYQCKV
RRV    YEHTATIPNVVGFPYKAHIERNGFSPMTLQLEVVETSWEPTLNLEYITCEYKTVVPSPFIKCCGTSECSSKEQPDYQCKV
SINDV  YEHATTVPNVPQIPYKALVERAGYAPLNLEITVMSSEVLPSTNQEYITCKFTTVVPSPKIKCCGSLECQPAAHADYTCKV
VEEV   YEHATTMPSQAGISYNTIVNRAGYAPLPISITPTKIKLIPTVNLEYVTCHYKTGMDSPAIKCCGSQECTPTYRPDEQCKV
EEEV   YEHTAVMPNKVGIPYKALVERPGYAPVHLQIQLVNTRIIPSTNLEYITCKYKTKVPSPVVKCCGATQCTSKPHPDYQCQV

                90       100       110       120       130       140       150
MAYV   FTGVYPFMWGGAYCFCDSENTQMSEAYVERADVCKHDYAAAYRAHTASLRAKIKVTYGTVN.QTVEAYVNGDHAVTIAGT
CHIKV  FTGVYPFMWGGAYCFCDAENTQLSEAHVEKSESCKTEFASAYRAHTASASAKLRVLYQGNN.ITVTAYANGDHAVTVKDA
SFV    YTGVYPFMWGGAYCFCDSENTQLSEAYVDRSDVCRHDHASAYKAHTASLKAKVRVMYGNVN.QTVDVYVNGDHAVTIGGT
RRV    YTGVYPFMWGGAYCFCDSENTQLSEAYVDRSDVCKHDHASAYKAHTASLKATIRISYGTIN.QTTEAFVNGEHAVNVGGS
SINDV  FGGVYPFMWGGAQCFCDSENSQMSEAYVELSADCASDHAQAIKVHTAAMKVGLRIVYGNTT.SFLDVYVNGVTPGTSKDL
VEEV   FTGVYPFMWGGAYCFCDTENTQVSKAYVMKSDDCLADHAEAYKAHTASVQAFLNITVGEHS.IVTTVYVNGETPVNFNGV
EEEV   FTGVYPFMWGGAYCFCDTENTQMSEAYVERSEECSIDHAKAYKVHTGTVQAMVNITYGSVTWRSADVYVNGETPAKIGDA

     160       170       180       190       200       210       220       230
MAYV   KFIFGPVSTAWTPFDTKIVVYKGEVYNQDFPPYGAGQPGRFGDIQSRTLDSKDLYANTGLKLARPAAGNIHVPYTQTPSG
CHIKV  KFIVGPMSSAWTPFDNKIVVYKGDVYNMDYPPFGAGRPGQFGDIQSRTPESKDVYANTQLVLQRPAAGTVHVPYSQAPSG
SFV    QFIFGPLSSAWTPFDNKIVVYKDEVFNQDFPPYGSGQPGRFGDIQSRTVESNDLYANTALKLARPSPGMVHVPYTQTPSG
RRV    KFIFGPISTAWSPFDNKIVVYKDDVYNQDFPPYGSGQPGRFGDIQSRTVESKDLYANTALKLSRPSPGVVHVPYTPTPSG
SINDV  KVIAGPISASFTPFDHKVVIHRGLVYNYDFPEYGAMKPGAFGDIQATSLTSKDLIASTDIRLLKPSAKNVHVPYTQASSG
VEEV   KITAGPLSTAWTPFDRKIVQYAGEIYNYDFPEYGAGQPGAFGDIQSRTVSSSDLYANTNLVLQRPKAGAIHVPYTQAPSG
EEEV   KLIIGPLSSAWSPFDNKVVVYGHEVYNYDFPEYGTGKAGSFGDLQSRTSTSNDLYANTNLKLQRPQAGIVHTPFTQAPSG

     240       250       260       270       280       290       300       310
MAYV   FKTWQKDRDSPLNAKAPFGCTIQTNPVRAMNCAVGNIPVSMDIADSAFTRLTDAPIISELLCTVSTCTHSSDFGGVAVLS
CHIKV  FKYWLKERGASLQHTAPFGCQIATNPVRAMNCAVGNMPISIDIPDAAFTRVVDAPSLTDMSCEVPACTHSSDFGGVAIIK
SFV    FKYWLKEKGTALNTKAPFGCQIKTNPVRAMNCAVGNIPVSMNLPDSAFTRIVEAPTIIDLTCTVATCTHSSDFGGVLTLT
RRV    FKYWLKEKGSSLNTKAPFGCKIKTNPVRAMDCAVGSIPVSMDIPDSAFTRVVDAPAVTDLSCQVVVCTHSSDFGGVATLS
SINDV  FEMWKNNSGRPLQETAPFGCKIAVNPLRAVDCSYGNIPISIDIPNAAFIRTSDAPLVSTVKCEVSECTYSADFGGMATLQ
VEEV   FEQWKKDKAPSLKFTAPFGCEIYTNPIRAENCAVGSIPLAFDIPDALFTRVSETPTLSAAECTLNECVYSSDFGGIATVK
EEEV   FERWKRDKGAPLNDVAPFGCSIALEPLRPENCAVGSIPISIDIPDAAFTRISETPTVSDLECKITECTYASDFGGIATLP

     320       330       340       350       360       370       380       390
MAYV   YKVEKAGRCDVHSHSNVAVLQE..VSIEAEGRSVIHFSTASAAPSFIVSVCSSRATCTAKCEPPKDHVVTYPANHNGITL
CHIKV  YAVSKKGKCAVHSMTNAVTIREAEIEVEGNSQLQISFSTALASAEFRVQVCSTQVHCAAECHPPKDHIVNYPASHTTLGV
SFV    YKTNKNGDCSVHSHSNVATLQEATAKVKTAGKVTLHFSTASASPSFVVSLCSARATCSASCEPPKDHIVPYAASHSNVVF
RRV    YKTDKPGKCAVHSHSNVATLQEATVDVKEDGKVTVHFSTASASPAFKVSVCDAKTTCTAACEPPKDHIVPYGASHNNQVF
SINDV  YVSDREGQCPVHSHSSTATLQESTVHVLEKGAVTVHFSTASPQANFIVSLCGKKTTCNAECKPPADHIVSTPHKNDQEFQ
VEEV   YSASKSGKCAVHVPSGTATLKEAAVELTEQGSATIHFSTANIHPEFRLQICTSYVTCKGDCHPPKDHIVTHPQYHAQTFT
EEEV   TNPVKQETVQFIVHQVLQLLKRMTSPLLRAGSFTFHFSTANIHPAFKLQVCTSGITCKGDCKPPKDHIVDYPAQHTESFT

         400       410       420       430
MAYV   PDLSSTAMTWAQHLAGGVGLLIALAVLILVIVTC..ITLRR..
CHIKV  QDISATAMSWVQKITGGVGLVVAVAALILIVVLC..VSFSRH.
SFV    PDMSGTALSWVQKISGGLGAFAIGAILVLVVVTC..IGLRR..
RRV    PDMSGTAMTWVQRLASGLGGLALIAVVVLVLVTC..ITMRR..
SINDV  AAISKTSWSWLFALFGGASSLLIIGLMIFACSMM..LTSTRR.
VEEV   AAVSKTAWTWLTSLLGGSAVIIIIGLVLATIVAMYVLTNQKHN
EEEV   SAISATAWSWLKVLVGGTSAFIVLGLIATAVVAL.VLFFHRH.
```

**C - Multiple Sequence Alignment of the Alphaviral E2**

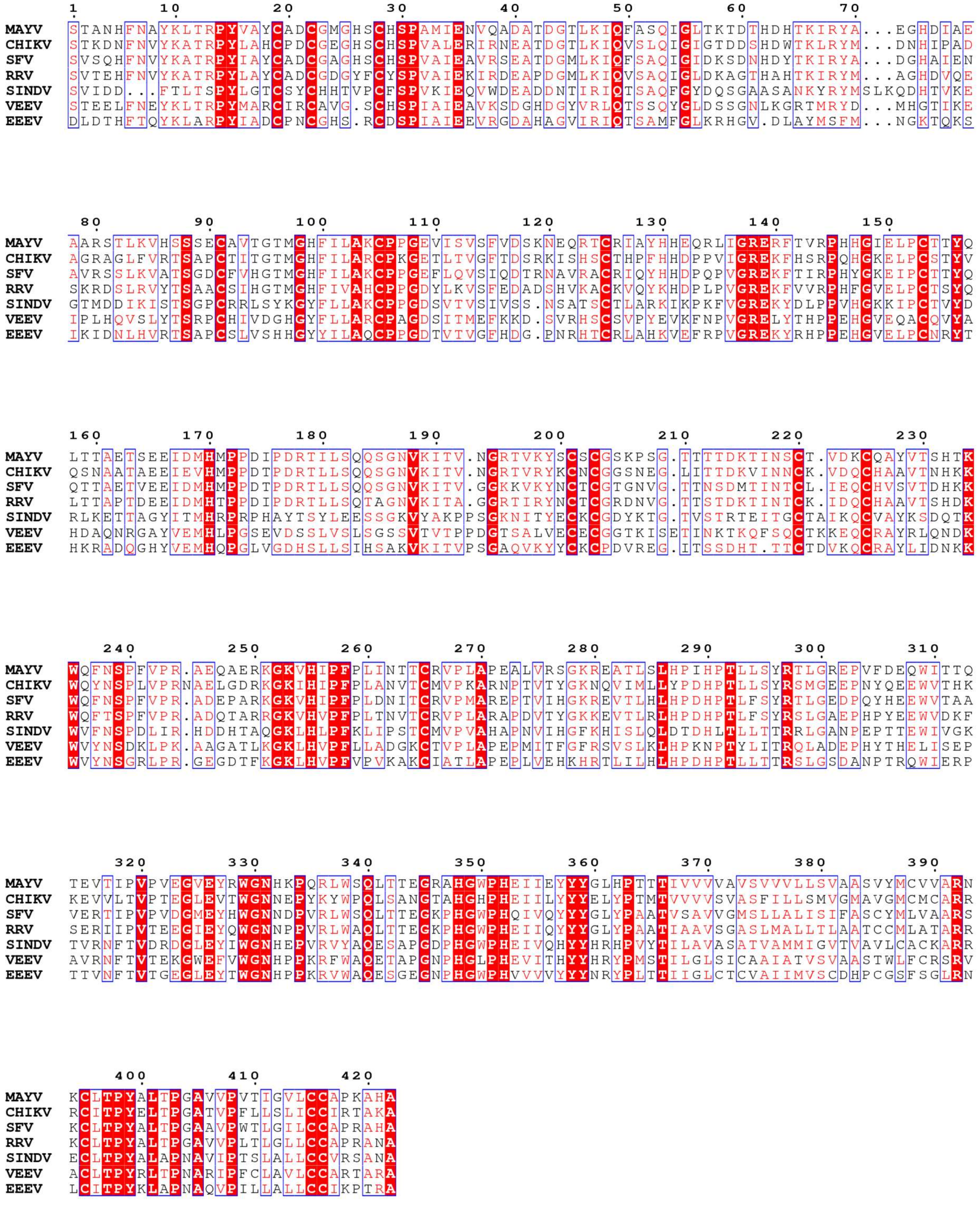

**Figure S4. Multiple sequence alignment of alphaviruses structural proteins.** The alignments were performed for proteins C (**A**), E1 (**B**) and E2 (**C**) using the Muscle Algorithm in EBI server (https://www.ebi.ac.uk/Tools/msa/muscle/) and visualized with ESPript version 3.0 server.

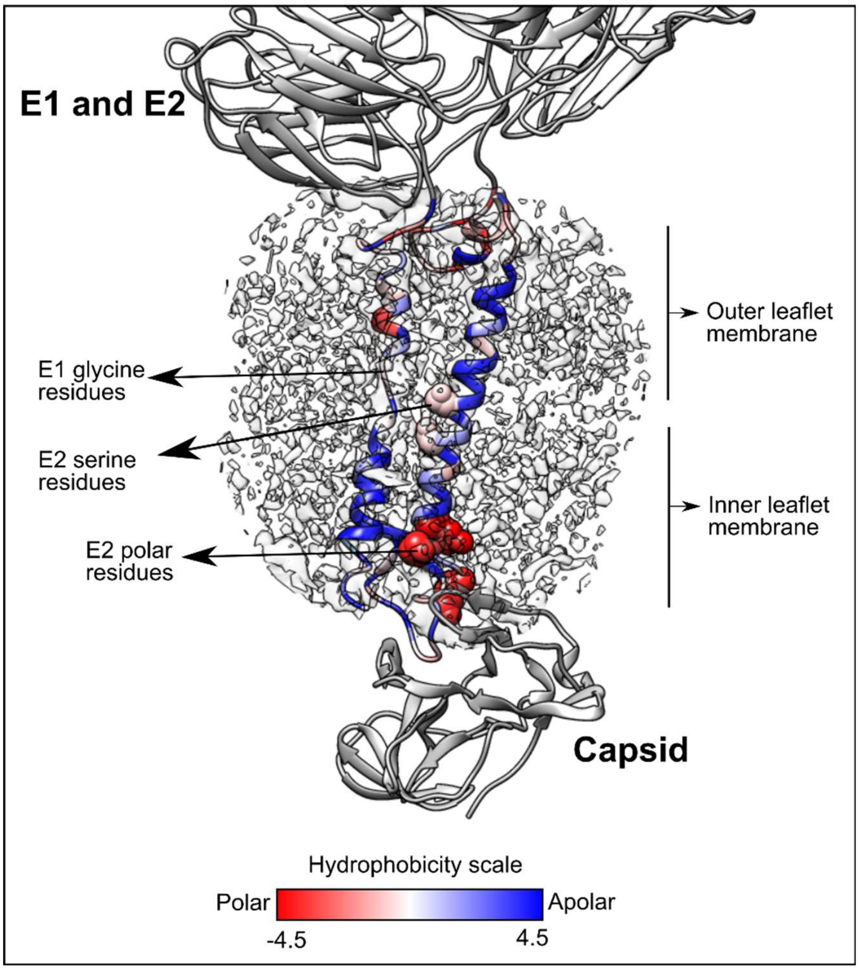

**Figure S5. MAYV E1 and E2 transmembrane domains.** Overall view of E1 and E2 TM domains inserted into the lipid membrane. The 3D atomic model is colored by hydrophobicity using Kyte-Doolittle scale. Important arginine and lysine residues in E2 are highlighted in spheres, as well as glycine (E1) and serine (E2) residues. The electron density is shown in grey surface.

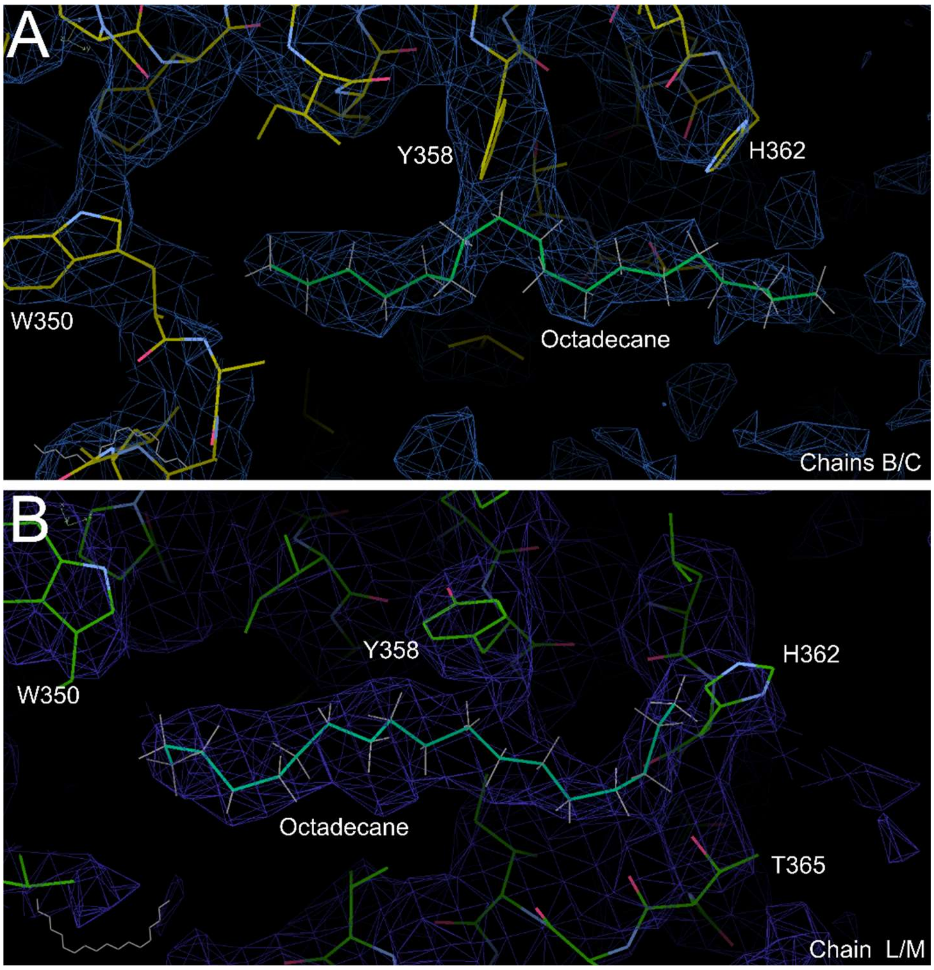

**Figure S6. 3D model fitting of a C18 hydrocarbon (Octadecane) in the extra density from two MAYV E1-E2 heterodimers (chains B/C and chains L/M).** Octadecane was built and fitted into the density map using Coot. The density map was rendered at 2.5 sigma contour level.

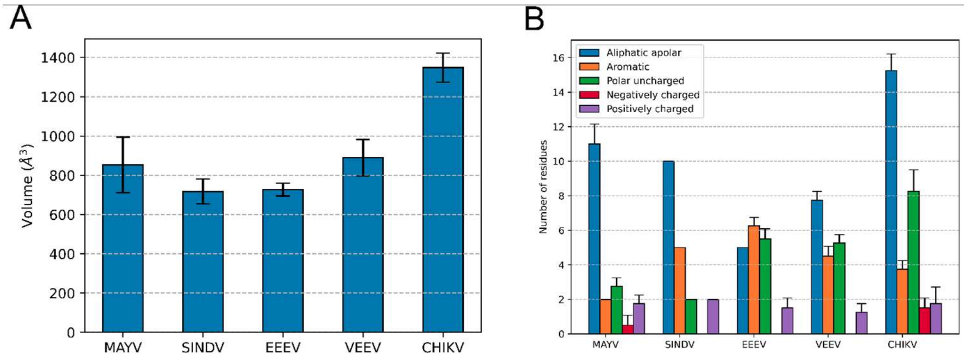

**Figure S7. Structural features extracted from the cavity between E1 and E2 TM helices in MAYV and other alphaviruses.** (A) Cavity volume estimated for the four E1-E2 heterodimers in asymmetric unit. (B) Number of residues in each four E1-E2 heterodimers in asymmetric unit separated by classes. Aliphatic apolar: ALA, VAL, ILE,

LEU, GLY, PRO; Aromatic: PHE, TYR, TRP; Polar uncharged: SER, THR, CYS, MET, ASN, GLN; Negatively charged: GLU, ASP; Positively charged: ARG, LYS, HIS.

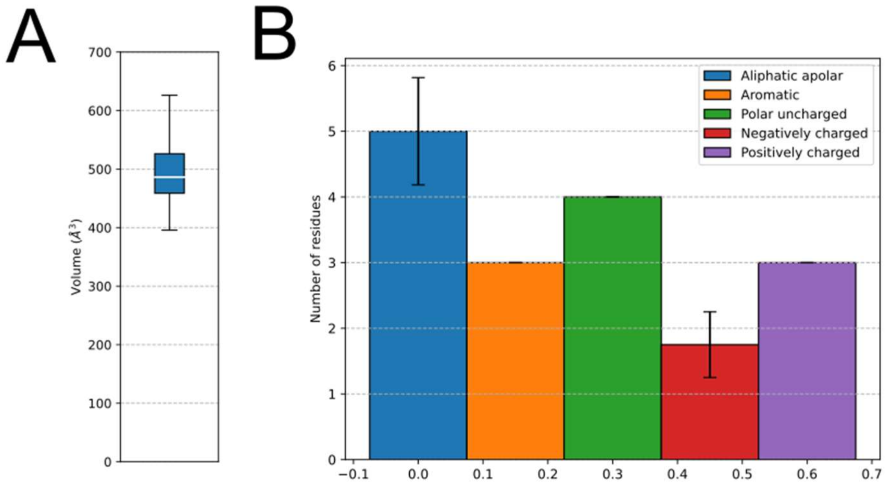

Figure S8. **Structural features extracted from the C-protein cavity that binds to E2 C-terminal.** (A) Boxplot of cavity volume estimated for the four capsids in asymmetric unit. (B) Number of residues in each four capsids in asymmetric unit separated by classes. Aliphatic apolar: ALA, VAL, ILE, LEU, GLY, PRO; Aromatic: PHE, TYR, TRP; Polar uncharged: SER, THR, CYS, MET, ASN, GLN; Negatively charged: GLU, ASP; Positively charged: ARG, LYS, HIS.

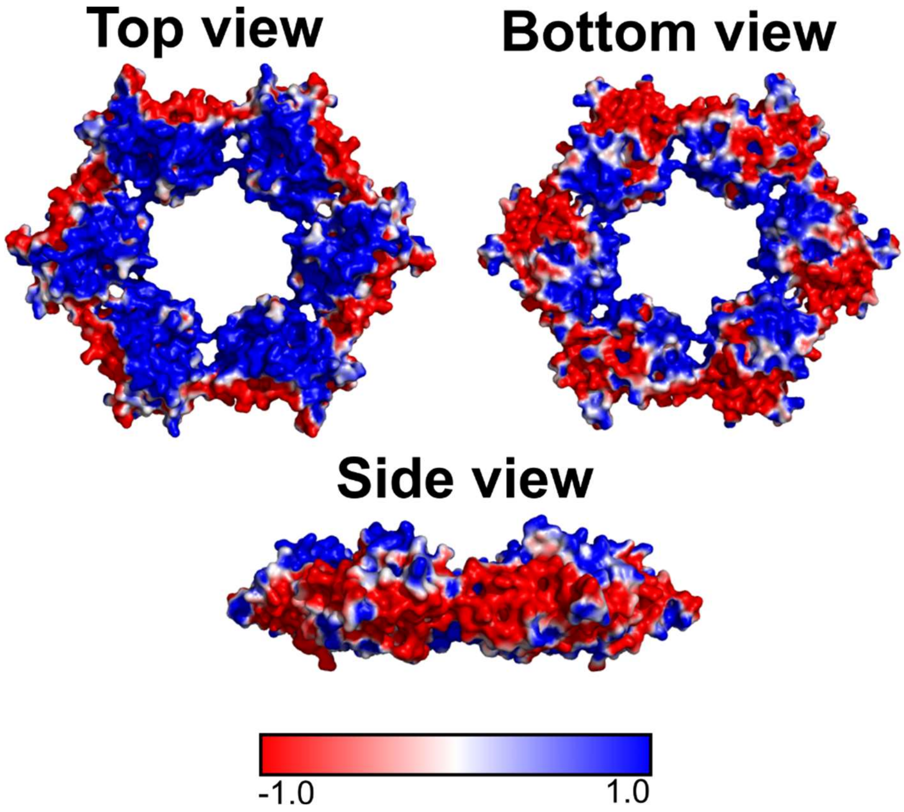

**Figure S9. The electrostatic potential of at the surface of MAYV capsid proteins.** The electrostatic potential was calculated using ABPS and visualized in Pymol. Top, bottom and side view from capsid hexameric organization is shown. Charges are presented in a gradient of blue (positive) to red (negative), white being neutral local charge.

**Table S1. Identity between structural proteins from MAYV and other alphaviruses.**

| | **CHIKV strain S27** | **SFV** | **RRV strain T48** | **SINDV** | **EEEV strain Florida 91-469** | **VEEV strain TC-83** |
|---|---|---|---|---|---|---|
| **Capsid – disorder N-terminal (1-102)** | **53 %** | **56 %** | **63 %** | **43 %** | **33 %** | **37 %** |
| **Capsid - Structured domain (103-258)** | **89 %** | **90 %** | **90 %** | **65 %** | **63 %** | **64 %** |
| **E1** | **62 %** | **72 %** | **71 %** | **48 %** | **52 %** | **52 %** |
| **E2** | **56 %** | **67 %** | **62 %** | **37 %** | **41 %** | **38 %** |

**Table S2. Cryo-EM data collection and processing.**

| | |
|---|---|
| **Symmetry imposed** | **Icosahedral** |
| **Number of micrographs** | **175840 (8792 movie stacks)** |
| **Initial number of particle images** | **79000** |
| **Number of particle images used in the final 3D reconstruction** | 40179 |
| **Map global resolution** | 4.4 (½ bit threshold) 4.3 (0.143 threshold) |

**Table S3. Overall MAYV 3D atomic model quality evaluated by MolProbity.**

| | |
|---|---|
| **Structural composition** | |
| **Chains** | **12** |
| **Residues** | **4004** |
| **MolProbity statistics** | |
| **Bonds length (Å) (RMSD - # > 4 sigmas)** | **0.006 (0)** |
| **Bond angles (º) (RMSD - # > 4 sigmas)** | **1.070 (43)** |
| **MolProbity score** | **2.0** |
| **Clash score** | **8.7** |
| **Ramachandran plot (%)** | |
| **Outliers** | **0.1** |
| **Allowed** | **11.4** |
| **Favored** | **88.6** |
| **Rotamer outliers (%)** | **0.59** |
| **Cβ outliers (%)** | **0** |
| **Model vs. Data** | |
| **CC (mask)** | **0.69** |

**Table S4. List of alphavirus PDB files and Cryo-EM maps used for comparative studies.**

| Alphavirus | PDB ID | EMD ID | Method | Resolution | Reference |
|---|---|---|---|---|---|
| CHIKV | 6NK5, 6NK6 | 9393 | Cryo-EM | ~4.5 A | Basore, 2019 |
| SINDV (only E1 and E2) | 6IMM | 9693 | Cryo-EM | ~3.5 A | Chen, 2018 |
| VEEV | 3J0C | 5275 | Cryo-EM | ~4.4 A | Zhang, 2011 |
| EEEV | 6MX4 | 9280 | Cryo-EM | ~4.4 A | Hasan, 2018 |

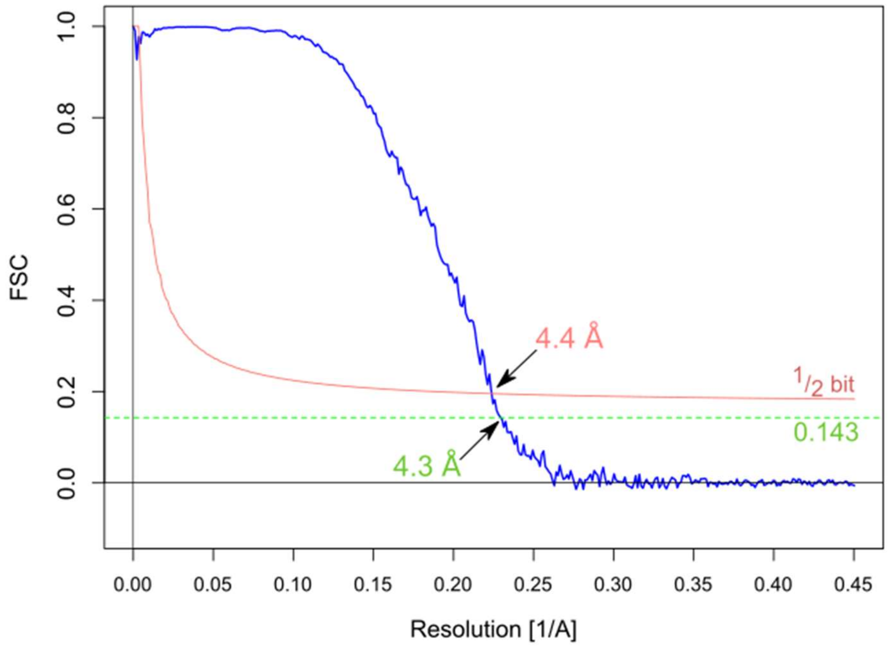

**Figure S10.** Fourier shell correlation profile of MAYV Cryo-EM density map. FSC curve calculated from two MAYV 3D volumes is shown in blue,the ½--bit curve in red and the 0.143 crossing line in green. The global resolution determined based on the ½--bit or 0.143 criterion is indicated[60].
